# Modelling the Dynamics of Biological Systems with the Geometric Hidden Markov Model

**DOI:** 10.1101/224063

**Authors:** Borislav Vangelov, Mauricio Barahona

## Abstract

Many biological processes can be described geometrically in a simple way: stem cell differentiation can be represented as a branching tree and cell division can be depicted as a cycle. In this paper we introduce the geometric hidden Markov model (GHMM), a dynamical model whose goal is to capture the low-dimensional characteristics of biological processes from multivariate time series data. The framework integrates a graph-theoretical algorithm for dimensionality reduction with a latent variable model for sequential data. We analyzed time series data generated by an in silico model of a biomolecular circuit, the represillator. The trained model has a simple structure: the latent Markov chain corresponds to a two-dimensional lattice. We show that the short-term and long-term predictions of the GHMM reflect the oscillatory behaviour of the genetic circuit. Analysis of the inferred model with a community detection methods leads to a coarse-grained representation of the process.

## Introduction

Biological processes such as disease progression, stem cell differentiation or the cellular response to external stimuli can be investigated using samples characterizing the cellular state at successive time periods^1^. Although each sample can measure a large number of variables (e.g. gene expression, protein abundances, metabolite levels), many processes can be represented in a simple way. The relation between the different stages of stem cell differentiation and of cell division are often depicted as a branching tree^2^ and as a cycle correspondingly. The simple structure of these processes can be represented in a low-dimensional space and therefore we conjecture that the dynamics of the cell state might be captured by a noisy low-dimensional manifold. In this paper we describe a framework that learns a low-dimensional dynamical model from multivariate time series data tracing the evolution of the process. With the proposed framework we infer an interpretable model that allows us to understand the structure of the process, the relation between different stages and to predict how the system state might evolve over time.

A low-dimensional latent representation of the data can be obtained both with model-based and with geometry-based approaches. Model-based approaches for dimensionality reduction estimate a model that maps from the high-dimensional space to a low-dimensional latent space (e.g. principal component analysis), from the low-dimensional space to the high-dimensional space (e.g. probabilistic principal component analysis^3^, Gaussian process latent variable model^4^) or by combining two models: one mapping from the high-dimensional space to the low-dimensional space and another from the low-dimensional space to the high-dimensional space (e.g. autoencoders). The properties of the specified model determine to a large extent the power of model-based methods to capture the complex geometry of high-dimensional datasets (e.g. a linear model can not recover the structure of a dataset on a nonlinearly embedded manifold). Geometry-based methods use a different approach for dimensionality reduction. They extract information about the geometry of the high-dimensional data and then project the samples so that the projected samples have similar geometric characteristics in the low-dimensional space. The geometry can be described by the pairwise distances between the samples (e.g., multidimensional scaling) or by a network linking samples that are local neighbours (e.g., Isomap^5^, LLE^6^). We are especially interested in the latter methods as the connectivity of the network can capture the continuity of nonlinear manifolds. Graph-theoretical methods have been previously applied to extract a low-dimensional representation of high-dimensional biomolecular data in order to understand the relationship and the similarity of different samples (different treatments, conditions or cell types)^7^.

The advantage of geometry-based methods is that they do not make specific assumptions about the mapping between the low-dimensional and the high-dimensional space and have simpler objective functions. However, geometry-based algorithms treat the observations as independent samples and even extensions designed for time series data ignore the directionality of time^8,9^, an important feature of time series data. Additionally, although they can recover the low-dimensional structure of the process they can not predict how the system state might evolve in time. Model-based methods have been extended to take into account the time order of the samples. Latent variable models for sequential data integrate a function describing the dynamics in the latent space and a function mapping from the latent space to the original high-dimensional space. A wide range of latent variable models for time series data have been proposed in the literature: the hidden Markov model (HMM)^10^, linear dynamical systems^11^, the Gaussian process dynamical model^12^. Latent variable models have been applied to model how T-cells^13^ and stem cells^14^ change over time.

In this paper, we introduce the geometric hidden Markov model (GHMM), a framework that combines a geometry-based and a model-based approach. The GHMM can be applied to multivariate time series data assuming that the observed trajectories lie on a low-dimensional manifold. Learning a GHMM model consists of three steps. The first step is the projection of the data to a low-dimensional space using the RMST-Isomap algorithm^15^, a modified version of the well known Isomap algorithm^5^. In the second step, Gaussian process regression (GPR)^16^ models mapping from the projected coordinates to each of the original dimensions are estimated. Finally, a HMM with a lattice structure is created, its prior beliefs are intialised using the GPRs and it’s trained using the variational Bayes approach^17^. Whereas previous works use the coordinates of the projected samples as direct input to the latent variale model^18–20^, we use them only to initialise the prior beliefs for the latent states and the training is still performed on the original data.

We illustrate the GHMM with trajectories generated by a stochastic model of the repressilator^21^, a biomolecular circuit exhibiting oscillatory behavior. We train a GHMM whose latent states can be represented as a two-dimensional lattice. We consider the predictions of the model in the short-term and the systems expected long-term behavior. In particular, we focus to what extent the trained model reflects the oscillatory behavior of the repressilator. Additionally, we show how a coarse-grained description of the dynamics can be extracted from the GHMM by partitioning the latent Markov chain in communities (i.e. latent states connected with high probabilities in the transition matrix). Although here we focus on the application of the GHMM to the analysis of data originating from biological processes, it can also be applied to domains where the dynamics of high-dimensional systems are described using the “landscape” paradigm (e.g. evolutionary processes^22^, protein folding^23^, molecular processes^24^).

## Results

We use data from an in silico model of the repressilator^21^ to showcase the GHMM. The repressilator consists of a sequence of genes where every gene represses the expression of the next gene in the sequence and the last gene represses the expression of the first gene (Fig. 1A). Repressilators with an odd number of genes exhibit oscillatory behavior. We use the following stochastic model to describe the four biochemical processes:

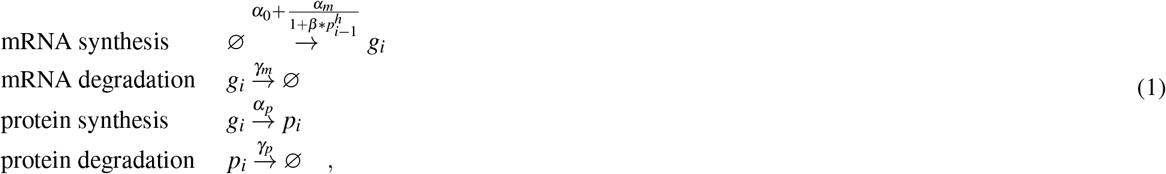

**Figure 1.**
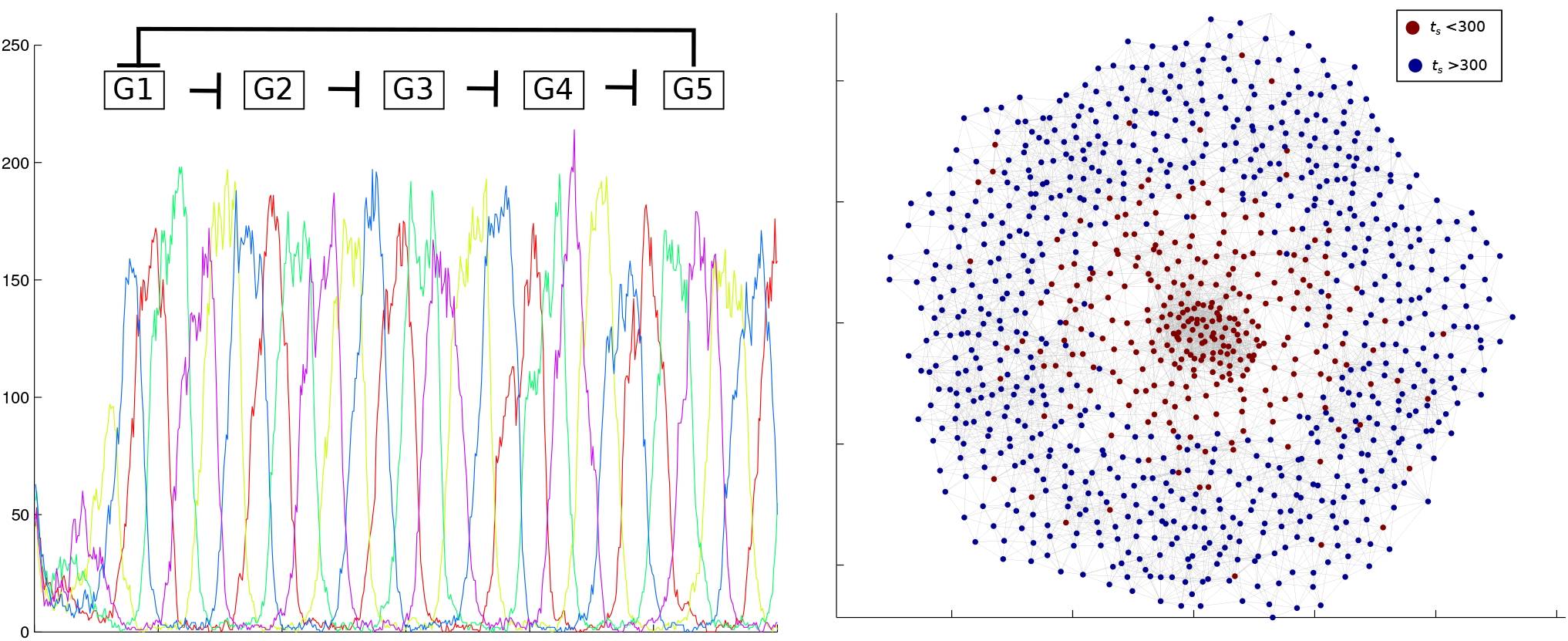
A) Single simulation of the mRNA profiles for a repressilator with 5 genes. B) Projection of the samples to a plane with the RMST-Isomap algorithm.

We simulated 100 trajectories with a repressilator with five genes using the Gillespie algorithm^25^, where the initial condition is no abundance of genes’ products. In Figure 1A, we show a single time trace with the five mRNA profiles. After an initial transient stage the model shows stable oscillatory behavior. From now on, we focus only the mRNA expression as often either the mRNA or the protein levels are observed in a single experiment.

The first step of the GHMM is the projection of the samples to a low-dimensional space with the RMST-Isomap algorithm. Due to the large number of samples we selected thousand samples with the following strategy: initially the samples were randomly chosen and then we updated the selection so as to maximize the minimal distance between the selected samples. In Figure 1B, we show the projected samples on the plane with the color-coding of the samples corresponding to the time component. Centrally located samples correspond to the initial time steps and samples corresponding to the later time steps form an orbit in the low-dimensional space. The second stage of the GHMM is the estimation of five GPR models mapping from the plane to each of the data dimensions. In Figure 2A we color-coded the samples with the expression profile of the first gene. We note that the gene expression profile has a well defined pattern in the low-dimensional space. The corresponding continuous GPR model captures the smooth changes in the gene expression levels. The GPR models for the other genes are shown in SI, Fig. 3,4.

**Figure 2.**
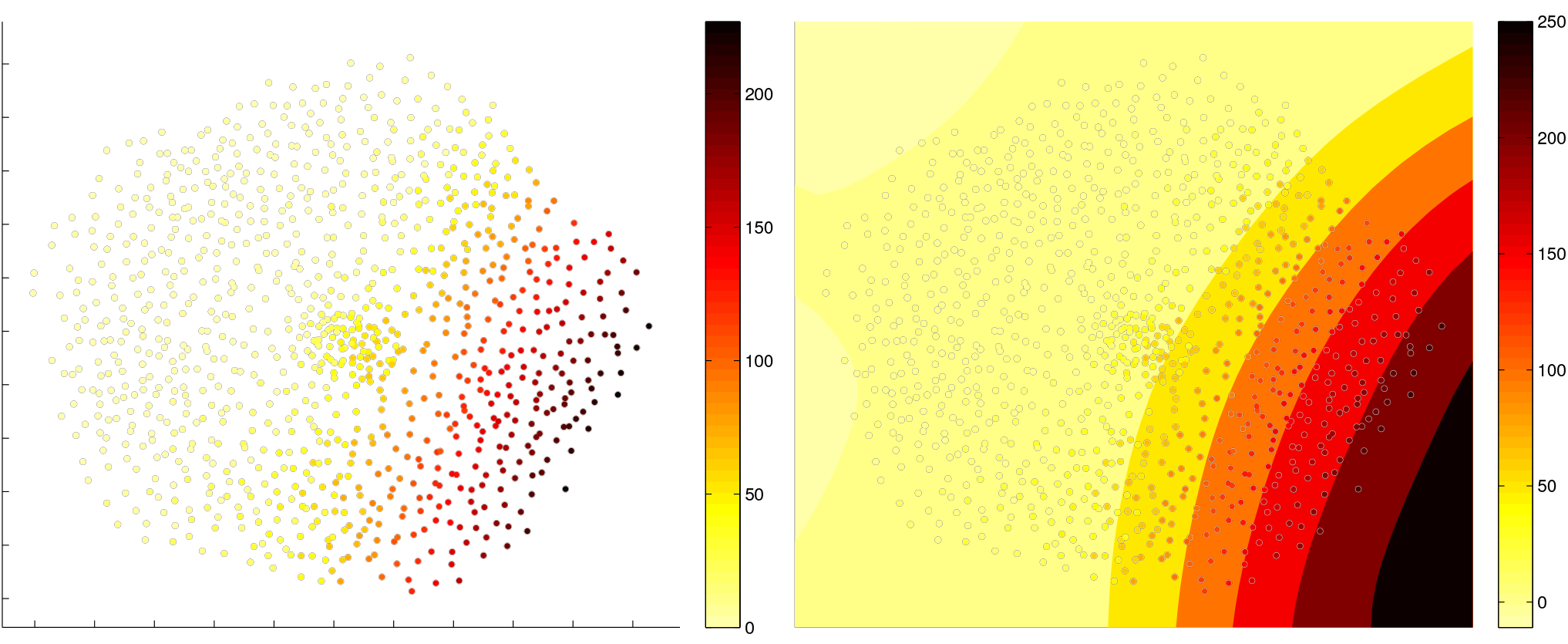
The color coding represents the abundance of the first proteins in the projected samples. A continuous model is estimated from the expression profile using the GPR.

Next, we initialized the prior beliefs for the latent variable model using the low-dimensional projection and the estimated regression models. We construct a lattice that serves as the prototype for the HMM. In the lattice every node is connected to at lost eight other nodes. Every node corresponds to a latent state and the prior beliefs for the emission probabilities are initialised using the GPRs. To find the optimal number of latent states we trained models with different number of hidden states and we explored how the variational free energy, a lower bound of the model evidence, changes (SI Fig.1). The variational free energy is maximal for a lattice of size 30 × 30. We note that although the latent Markov chain has a large number of latent states only a subset of them generate the observed samples with a significant probability (427 states are “responsible” from overall 900, SI Fig. 5). Further, we studied whether prior beliefs of a denser latent Markov chain can better account for the data. We trained models with different degree of connectivity and found that the variational free energy has a maximal value when a node in the lattice is connected to at most eight other nodes (see SI Fig. 2). An increase of the connectivity of the latent Markov chain results in a decrease of the model evidence. We believe that the prior beliefs of a sparse Markov chain can better account for the smooth change of the system state over time.

We used the trained latent Markov chain to predict the short-term dynamics of the system. In Figure 3 we show how the probability mass concentrated initially in a single latent state diffuses throughout the Markov chain after 10 and 40 periods. We note that the probability does not diffuse uniformly, but rather the diffusion follows a circular trajectory. We also looked at the inferred probability distribution in the first time period and at the stationary distribution of the latent Markov chain, i.e. in the limit when time is close to infinity. The probability mass in the first time period is concentrated in a few centrally located latent states as all stochastic simulations were initialized with the same initial conditions (SI Fig. 6A). The latent Markov chain is ergodic and has a unique stationary distribution. The stationary distribution informs about the expected long-term state of the system (SI Fig. 6B). The stationary distribution is concentrated in latent states with a circular arrangement capturing the oscillatory behavior of the system.

**Figure 3.**
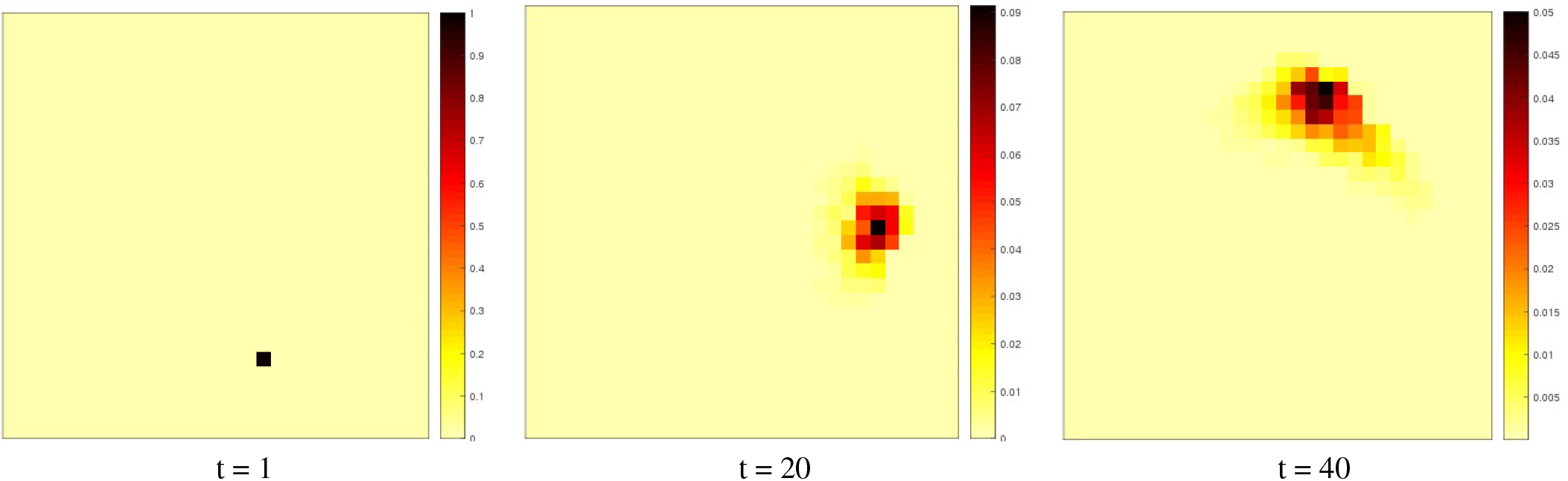
The diffusion of probability in the latent Markov chain shows the expected behavior of the system.

To understand the inferred latent Markov chain better, we looked also for a coarse-grained model of the dynamics. Here, we focused only on latent states “responsible” for the observed data. Latent states having a negligible likelihood of generating a sample are removed from the Markov chain and both the probability distribution in the first time period and the stochastic transition matrix are normalized. We employed Markov stability for community detection^26^ to find a coarse-grained representation of the Markov chain. Markov stability for community detection looks for partitions of the Markov chain with a well defined community structure. In a well defined community structure, a random walker is trapped in a particular community for a given Markov time. Markov time serves also as a resolution parameter, i.e. finer partitions are favored at small Markov times and coarse partitions are favored at larger Markov times. We looked for the optimal partitions with the Louvain algorithm^27^. We found robust partitions by comparing the ensemble of solutions obtained with the Louvain algorithm using the variation of information^28^.

The minimal variation of information was found at Markov time *M_t_* = 8 (SI Fig. 7). The optimal partition has six clusters as shown in Figure 4A. One of the clusters is located centrally where the probability mass in the initial time periods is concentrated. To understand the relation between the different clusters and the repressilator we estimated the number of latent states describing an overexpressed or an underexpressed gene. A gene is overexpressed in a latent state if the mean value of the inferred emission probability is greater than the average. The pattern of overexpressed/underexpressed genes in a cluster is shown in Figure 4B. We note that in the first cluster all genes have low expression values. In all the other five states, one gene is over-expressed, two genes are not expressed and two genes sequentially ordered in the circuit are expressed only partially. The arrows in the figure show how the probability mass diffuses in the coarse-grained Markov chain. The flow of probability follows a circulatory direction further reflecting the oscillatory behavior of the system.

**Figure 4.**
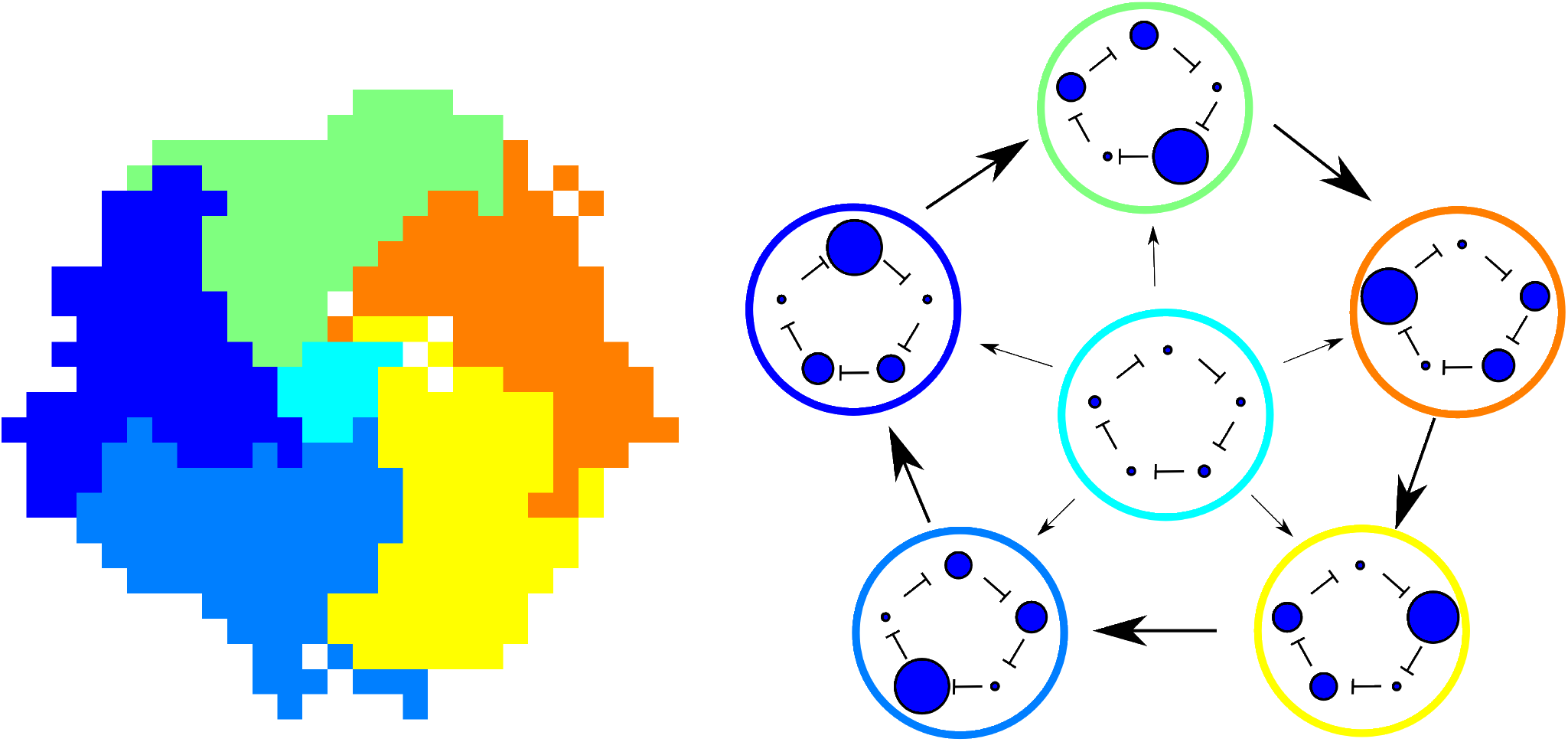
The inferred latent Markov chain can be approximated by a coarse-grained Markov chain with six states. Every state corresponds to a specific pattern of gene expression. Five of the states correspond to a pattern in which one gene is overexpressed, two are partially overexpressed and two are downregulated. The transition probabilities in the coarse-grained Markov chain reflect the oscillatory behavior of the system.

## Discussion

Multivariate time series data tracing the evolution of biological processes in time contains valuable information about the dynamics of biological systems. As we often assume that many processes have simple structure different techniques for dimensionality reduction have been widely applied to extract a low-dimensional representation of high-dimensional samples. These low-dimensional representations are a valuable resource that informs about the similarity between different samples corresponding to different cell types, different treatments or control vs. disease models. In the case of time series data the low-dimensional projection can reveal the relation between different stages of the process or how a biological system evolves under different conditions. However, the low-dimensional projection provides only a static picture of the process and ignores the factor of time contained in time series data. In this paper, we showed how the low-dimensional projection can be used also as a basis to learn a model of the system dynamics.

The GHMM extracts a dynamical model from time series data lying on a manifold embedded nonlinearly in a highdimensional space. The proposed methodology integrates a graph-theoretical algorithm for manifold learning with a latent variable model for sequential data. By integrating two different type of approaches, the GHMM can benefit from the advantages of both geometry-based and model-based algorithms. Firstly, the samples are projected nonlinearly to a low-dimensional space using a geometry-based algorithm for dimensionality reduction. The geometry-based method is computationally efficient and employs a simple optimization procedure. Most importantly, it focuses on the continuity of the data on the manifold and does not make any specific assumptions about the functional relation between the low-dimensional and the high-dimensional space.

In the second stage, we learn a latent variable model using prior beliefs derived from the low-dimensional projection. Whereas the low-dimensional projection informs only about the similarity between different samples, the latent variable model can capture also the dynamics of the system. Thus, it can predict how the systems state might evolve in the short term and what is the expected system state in the long term. To bridge the two approaches for data modeling we employ the low-dimensional projection as a basis and encode specific prior beliefs for each latent state. Firstly, we can encode our beliefs that the latent Markov chain has a sparse local connectivity due to the smoothness of the dynamical system. Prior belief of a sparse transition matrix allow us to efficiently learn large latent variable models. Secondly, we can encode our prior beliefs about the mapping between latent states and the observed data so that the emission probabilities trace the geometry of the low-dimensional manifold.

The procedure was applied on data generated with a biochemical stochastic model that exhibits oscillatory behavior. The projected samples reflect the continuity of the manifold: the coordinates of the original dimensions change smoothly in the low-dimensional space. Additionally, the optimal model has a sparse latent Markov chain corresponding to small transitions on the manifold in the high-dimensional space. The diffusion of probability in the latent Markov chain is non-uniform and follows a circular trajectory. The stationary probability distribution is concentrated in states with a circular arrangement on the lattice. The latter two observations show how the model captures the oscillatory behaviour of the repressilator. Finally, we applied a community detection method to extract a coarse-grained description of the model. By analyzing the diffusion of probability we found that there are six communities with latent states representing distinct gene expression patterns. Whereas the latent Markov chain approximates the continuity of the low-dimensional manifold, the model with six coarse states captures main features of the process.

We exemplified our approach with data generated from an in silico model of a biochemical circuit. In the future we plan to apply the methodology to experimental data characterizing biological processes such as the cell cycle and stem cell differentiation. We envisage that the approach can be applied also to the analysis of protein folding dynamics and evolutionary processes. Protein folding dynamics has been investigated using molecular dynamics simulations that generate high-dimensional trajectories of molecular conformations. Such trajectories have been analyzed with geometry-based for manifold learning in order to extract a low-representation of the protein folding landscape. The GHMM can provide further details on how the system evolves from one conformational state to other and what is the probability distribution of different states in the long term.

## Methods

Training the GHMM consists of three steps. The first step is the projection of the observed data to a low-dimensional space. We use the RMST-Isomap algorithm^15^, an extension of the well known Isomap algorithm for manifold learning. The second step is the estimation of regression models mapping from the coordinates of the projected samples to each of the original dimensions. The final step is the creation and the training of a HMM. The basis of the HMM is a two-dimensional lattice with sparse connectivity, where every node corresponds to a latent state and the connectivity is used as a prior for the latent Markov chain. The prior beliefs for the probabilistic models mapping from the latent states to the observed data are initialized using the regressions estimated in the previous step. The model is trained using a Variational Bayes approach^29^. Each of the steps is described in more details in the following subsections.

## Manifold Learning

Graph-theoretical methods for manifold learning capture the structure of a manifold by constructing a network in which a node represents an observation in the dataset and two nodes are connected in the network if the corresponding observations are local neighbors. The network models the dataset geometry and its connectivity traces the manifold continuity. The classical strategy for constructing a network from vector data is to link observations if the distance between them is below a threshold value or to connect every observation to its k-nearest neighbors. If the points are homogeneously distributed on the manifold, these two strategies can estimate a network structure that captures the geometry of the underlying manifold. However, if the data is not homogeneously distributed on the manifold, the estimated network will fail to represent the geometry correctly. If the threshold is small, finely sampled parts will be correctly described but the network will consist of several disconnected components. If the threshold is large, the network will be connected but well sampled parts of the manifold will be densely connected.

Thus we employ the RMST algorithm for network construction, an approach that is robust to in-homogeneously distributed data. The RMST expands a minimum spanning tree (MST), i.e. a spanning tree with minimal sum of edge weights, with a simple heuristic. The maximal edge weight in the MST path and the weight of the direct edge between two nodes are compared. If the maximal edge weight in the MST path is significantly smaller than the direct edge, then the MST path is a better model of the relation between the two points in the dataset as the path is a sequence of relatively small transitions. If the maximal edge weight in the MST path and the direct edge weight do not differ significantly, then there is no enough evidence that the MST path is a better model of the data continuity and the MST can be expanded by adding an edge between the two nodes. Let ***Y*** be a matrix with the observed data (***Y*** = [***y**_i_*,…,***y***_N_]), where each observation is a *D*-dimensional vector (***y**_i_* ∈ ℝ^d^). We compute a matrix with the pairwise distances between all vectors (*d_i,j_* = *d*(***y**_i_*, ***y**_j_*)) and we then construct a minimum spanning tree from the distance matrix. We apply the following heuristic to estimate the adjacency matrix ***E*** of the RMST:

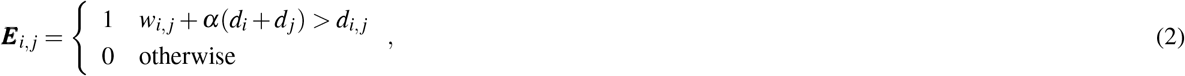

where *w_i,j_* is the maximal edge weight in the MST path between the ith and the jth node, α is a positive constant, *d_i_* is the distance to the *k* nearest neighbor of node *i*. The term *αd_i_* approximates the local data density around the ith observation and it is motivated by the Perturbed Minimum Spanning Tree algorithm^30^. The described heuristic combines both global characteristics of the data, the maximal edge weight in the MST path, and local characteristics, a term approximating the local data density around a sample.

The Isomap algorithm uses the network to define a new distance measure between the samples. The distance between two samples is the geodesic distance on the network between the corresponding nodes, i.e. the length of the shortest path. Finally, the samples are projected using the MDS algorithm given the matrix with pairwise distances between all samples. In this paper, we minimize the relative stress defined as:

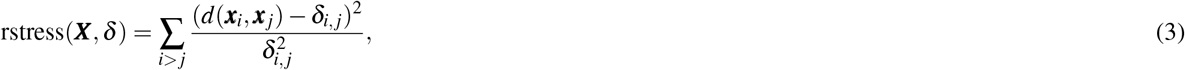

where ***x**_i_* is the projection on the low-dimensional space 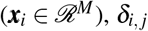 is the geodesic distance between the *i*th and the *j*th nodes on the network.

## Hidden Markov Models (HMM)

Here, we briefly review the HMM, one of the most popular latent variable models for sequential data. The HMM uses discrete latent states and the dynamics in the latent space is represented by a Markov chain. The relation between the latent states and the original data are described by the emission probabilities of every latent state. The state of a HMM with *K* latent states can be encoded with a latent variable *Z*, where *t* is the time. We use 1-of-K representation for the latent variable, i.e. ***z**^t^* is a ***K*** dimensional vector in which one of the elements is equal to one. The probability that the trajectory starts at a specific latent state is defined by the multinomial distribution 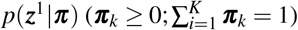. We use the Dirichlet distribution to define the prior for the multinomial distribution *p*(***π***) = Dir(***π***|***u***^π^). The latent dynamics are described by a Markov chain with transition matrix ***A***, where ***A**_i,j_* is the probability of transition from the *i*th to the *j*th state 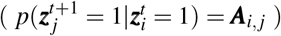. The *i*th row in the transition matrix (***A**_i_*) specifies the probabilities of transition from the ith latent state to the other latent states. The transition probabilities from a given state have multinomial distribution and we use the Dirichlet distribution *p*(***A**_i_*) = Dir(***A**_i_*|***u**^A^*) to model their prior. We use the multivariate Gaussian distribution to model the emission probabilities of the latent states. The multivariate Gaussian distribution of the ith state is defined by a mean ***μ**_i_* and a precision matrix **Λ**_*i*_. We employ the Gaussian-Wishart distribution to model the prior of the multivariate Gaussian distribution: 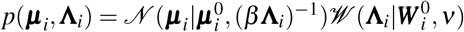.

The HMM can be inferred from data using the Variational Bayes (VB) framework^31^, a method for approximate Bayesian inference. The VB framework approximates the posterior distribution *P*(***Z***, ***θ***|***Y***) with a distribution *q*(***Z***, ***θ***) that can be factorized as:

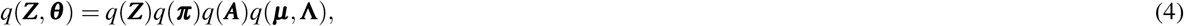

where ***Z*** is a matrix with the latent variables for all periods (***Z*** = [***z***^1^,…, ***z**^T^*]) and all model parameters are denoted by **θ** (**θ** = {***π**, **A***, ***μ***_1_, **Λ**_1_,…}). The VB framework decomposes the marginal likelihood of the data into the sum of the variational free energy and the Kullback-Leibler divergence between the approximate and the posterior distribution:

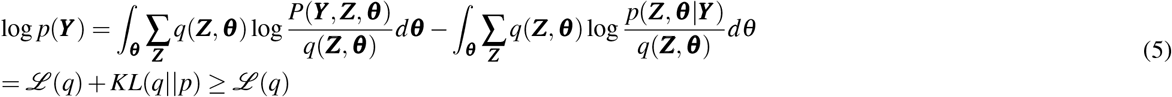

We infer a model by maximizing the variational free energy 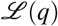. It is a lower bound of the marginal log-likelihood and can be used also as criterion for model selection, e.g. for finding the optimal number of latent states.

## Initialisation of the Prior Beliefs

The GHMM uses the low-dimensional projection obtained with the RMST-Isomap algorithm to initialize the prior beliefs for the parameters of the HMM. In the low-dimensional space we construct a lattice with sparse connectivity that serves as a basis for the latent Markov chain. We denote the coordinates of the lattice nodes by ***X***^⋆^ and the adjacency matrix of the lattice by ***L***. Every node in the lattice is a prototype of a latent state in the HMM. The sparse connectivity of the lattice encodes our prior beliefs of a sparse latent Markov chain with a non-zero probability of transition from a latent state only to its local neighbors. Our prior of a sparse latent Markov chain is due to the assumption that the time series data is generated by a continuous dynamical system and in a short period of time we expect only a small transition on the manifold. The prior probability distribution of the transitions from the *i*th state are initialized by *p*(***A**_i_*) = Dir(***A**_i_*|*u^L^**L**_i_*). The sparse latent Markov chain allows us to train efficiently large HMM. The prior beliefs for the first time period is that all latent states are equally likely (*p*(***π***) = Dir(***π***|*u^π^***1**)).

We use the low-dimensional projection to initialize also the priors for the emission models mapping from the latent states to the observed data. They are encoded using Gaussian-Wishart distributions for every latent state and we initialize them with different hyperparameters 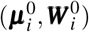. These hyperparameters are estimated with regression models that describe the relation between the projected data (***X***) and every dimension of the original data. The original samples are the *N* columns of the matrix 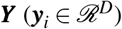 and a row in ***Y*** represents the coordinates of all *N* samples in a specific dimension 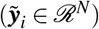. We use the Gaussian process regression (GPR) to infer a model between the independent variable, the coordinates of the projected samples ***X***, and the dependent variable, a single dimension of the original observations 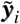. We assume that the dependent variables are a noisy realization of the true dependent variable where the noise comes from a Gaussian distribution with variance 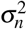. We estimate *D* independent regression models, one for each dimension of the original data. In the GPR, the observed data is modeled as a realization of a Gaussian process with a mean function (*μ*) and a kernel function (*k*). Here, we use the squared exponential kernel function and a constant mean function equal to zero. The GPR estimates a normal distribution that maps from a point on the lattice ***x**_k_* to the ith dimension of the high dimensional space:

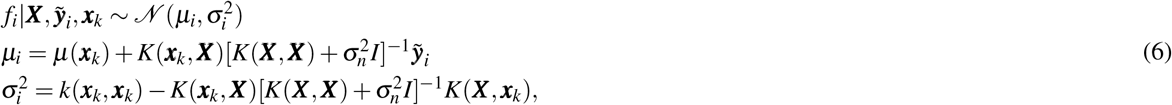

where 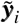 is a vector with the values of the projected samples in the *i*th dimension and the vector *K*(***X***, ***x**_k_*) contains the output of the kernel function applied on the pairwise combinations of ***X*** and ***x**_k_*. The hyperparameters of the models: the parameters of the kernel function, i.e. the length scale and noise variance, and the variance for the noise term 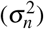 are computed by maximization of the marginal likelihood. We estimate *D* regression models, one for each dimension of the observed data. For every latent state, we combine the estimates of the GPR models to initialize the parameters of the Gaussian-Wishart distribution as 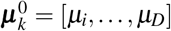 and 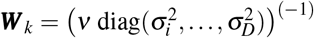. The parameters *ν* and *β* are constants and are equal for all states. As a result the prior beliefs for the emission probabilities of every latent state are different and they trace the geometry of the manifold embedded.

